## Supplementary for "Improved analyses of GWAS summary statistics by reducing data heterogeneity and errors"

Chen et al.

**Supplementary Figures 1-15**

**Supplementary Tables 1-16**

**Supplementary Notes**

### Supplementary Figures

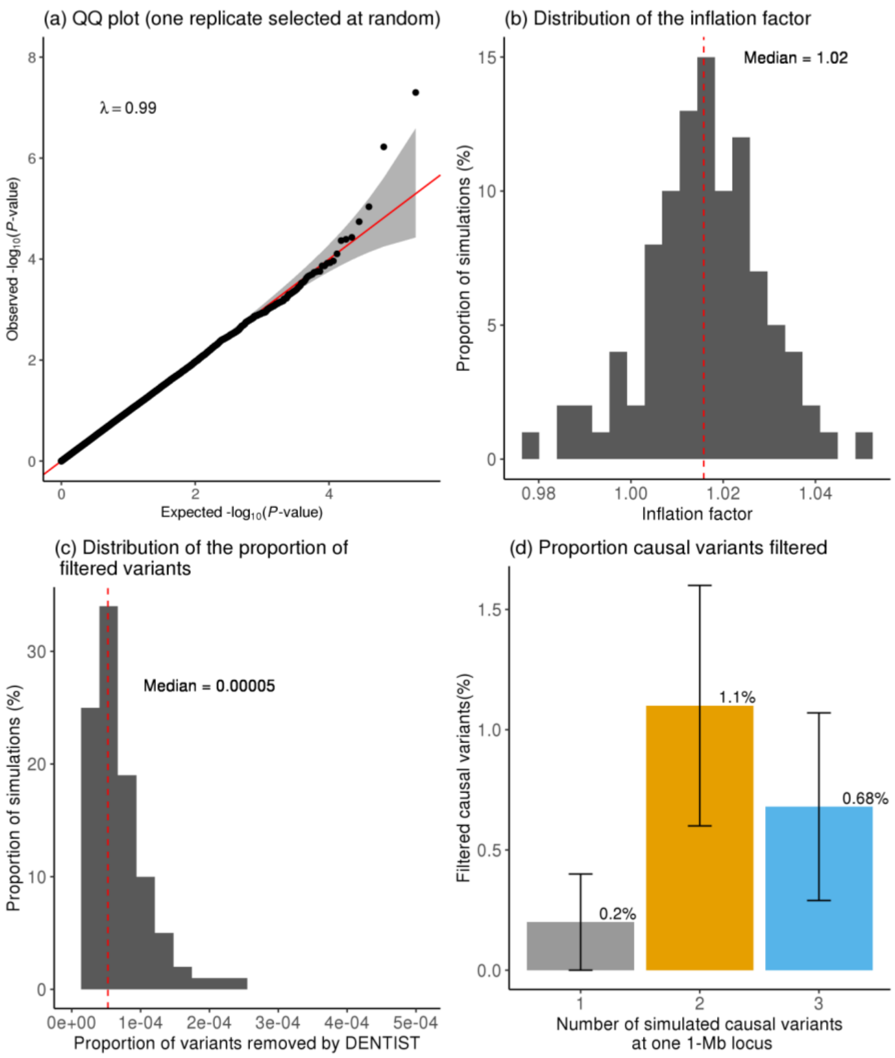


**Supplementary Figure 1. Statistical properties of the DENTIST test-statistic in the absence of errors.** We simulated a quantitative trait with 50 causal variants selected at random using the UK10K-WGS data without introducing genotyping or allelic errors. The inflation factor is defined as the median of the DENTIST test statistics of all the variants divided by 0.455, i.e., the expected median of $\chi_1^2$ under the null. The QQ plot of a simulation replicate selected randomly is shown in panel a. In panel b or c, the red vertical dashed line represents the median of the distribution. In panel (d), we performed additional simulations with one, two, or three causal variants ($r^2 < 0.1$ between the causal variants and $q^2 = 2\%$ for each causal variant) at a 1-Mb locus using the same data. Each scenario was repeated 440 times with the causal variants resampled in each replicate. Shown on the y-axis is the proportion of causal variants filtered by DENTIST across simulation replicates.

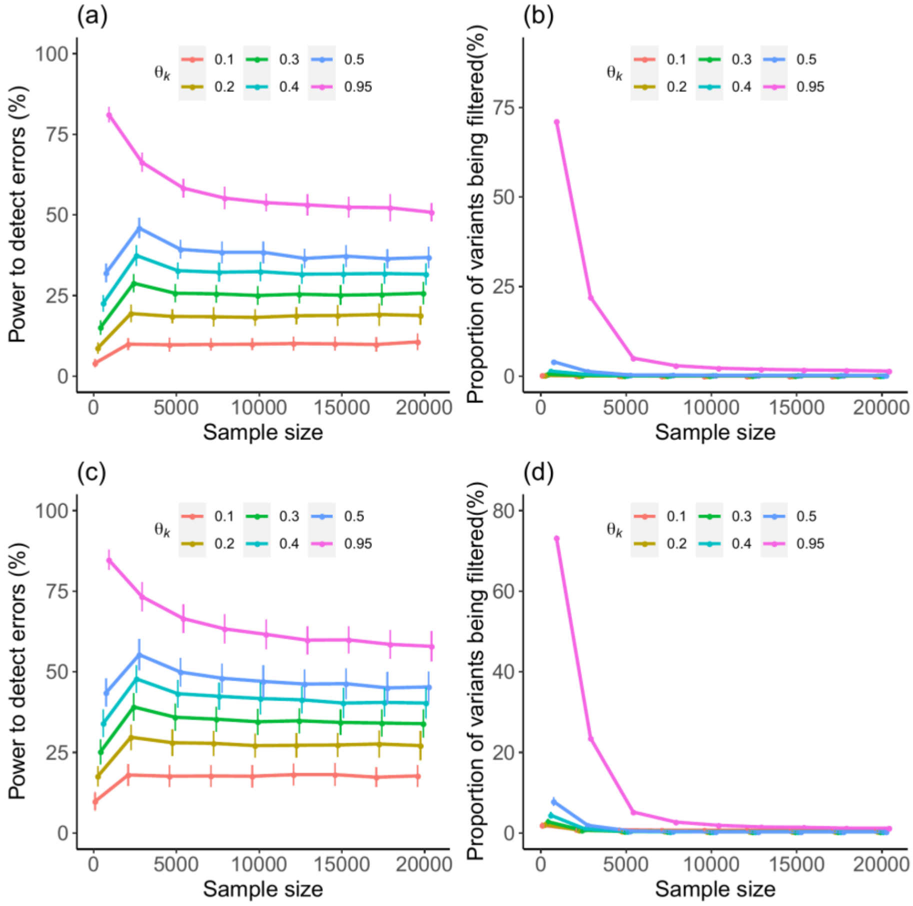


**Supplementary Figure 2. The choice of the SVD parameter $\theta_k$ and the reference sample size.** To investigate the effects of the SVD parameter $\theta_k$ (see Methods for the definition of $\theta_k$) and reference sample size on the performance of DENTIST, we analyzed the GWAS summary data simulated based on UK10K-WGS (n = 3,642) as described in the main text (simulating genotyping and allelic errors on 0.5% of the variants, respectively, on chromosome 22). UKBv3-20K (20,000 unrelated individuals from the UKB) was used as the LD reference for DENTIST analysis. Power is defined as the number of correctly identified erroneous variants by DENTIST divided by the total number of simulated erroneous variants. We repeated the analysis using a subset of the UKBv-20K data ($n_{\mathrm{ref}}$ = 500, 2,500, 5,000, 7,500, 10,000, 12,500, 15,000 or 17,500) with different levels of $\theta_k$ ($\theta_k = 0.1, 0.2, 0.3, 0.4, 0.5$ or 1). The results shown in panels a) and b) suggest the use $\theta_k = 0.5$ and $n_{\mathrm{ref}}$ > 5,000 in practice. We further performed simulations based on the genotype data of 20,000 unrelated UKB individuals (independent of UKBv3-20K) using the same strategy as in the UK10K simulation and using reference LD calculated from each subset of the UKBv-20K data ($n_{\mathrm{ref}}$ = 500, 2,500, 5,000, 7,500, 10,000, 12,500, 15,000 or 17,500). The results (panels c and d) show that the choice of $\theta_k$ and minimal $n_{\mathrm{ref}}$ do not seem to depend on discovery GWAS sample size.

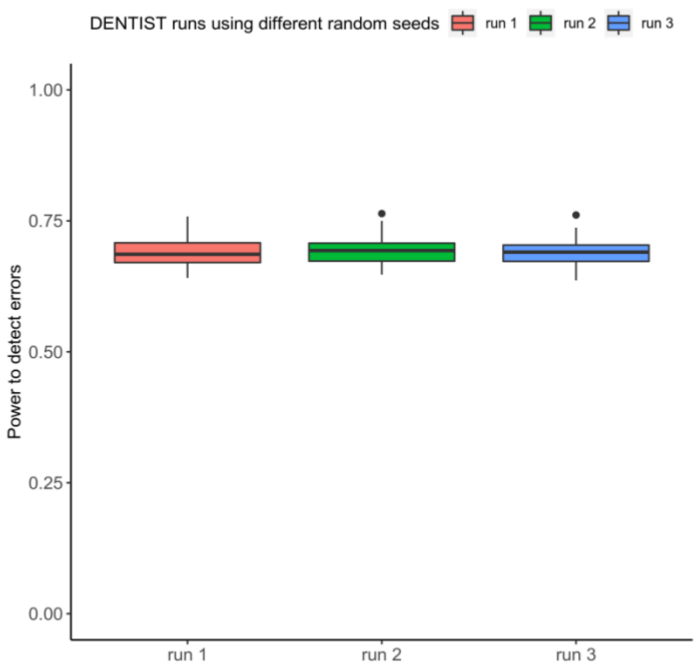


**Supplementary Figure 3. The stability of DENTIST to random seed.** We quantify how random seed used in DENTIST can affect the detection of errors by apply 3 DENTIST runs with different random seeds to GWAS summary data simulated based on UK10K-WGS (n = 3,642) as described in the main text (simulating genotyping and allelic errors on 0.5% of the variants, respectively, on chromosome 22). The power is quantified as the proportion of both errors detected from DENTIST based on 100 replicates.

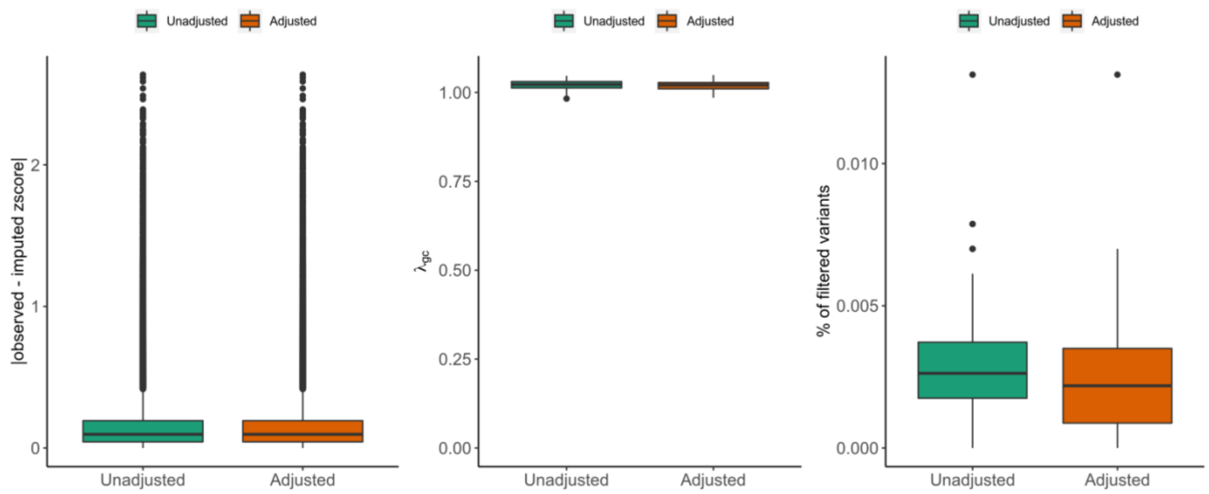


**Supplementary Figure 4. The impact of covariate adjustment on DENTST test.** Whether the GWAS summary statistics are from analysis with or without covariate adjustment is a source of heterogeneity between GWAS and LD reference because LD data are often computed from the reference without covariate adjustment. We used the UK10K-WGS data to simulate a polygenic trait (10,000 causal variants explaining 20% of the phenotypic variance in total) with 5% of the variance explained by the first two SNP-derived principal components (PCs). The simulation was repeated 100 times with the causal variants resampled in each replicate. Shown are comparisons of the DENTIST test statistics computed based on GWAS data adjusted for the first two PCs with those without PC correction.

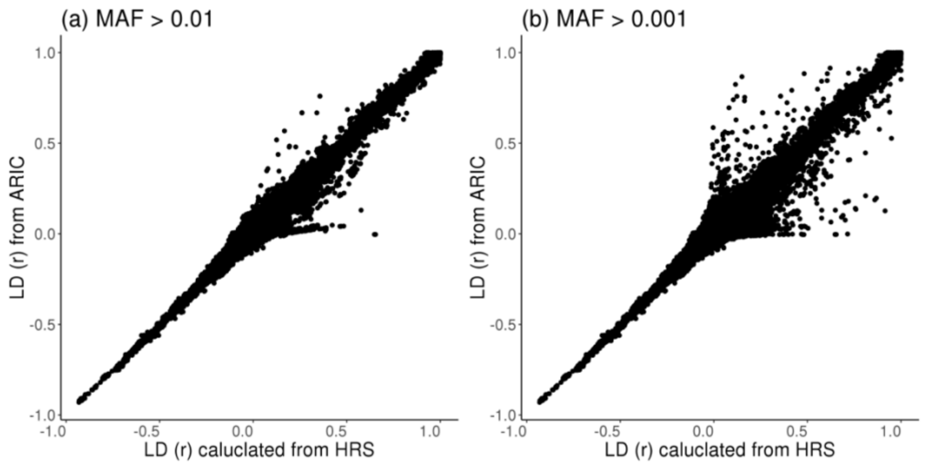


**Supplementary Figure 5. Comparison of LD between two samples of European ancestry.** Shown are LD correlations (r) between pairwise variants at a randomly selected locus on chromosome 22 computed from ARIC plotted against those from HRS. Panel a: variants with MAF >0.01. Panel b: variants with MAF >0.001.

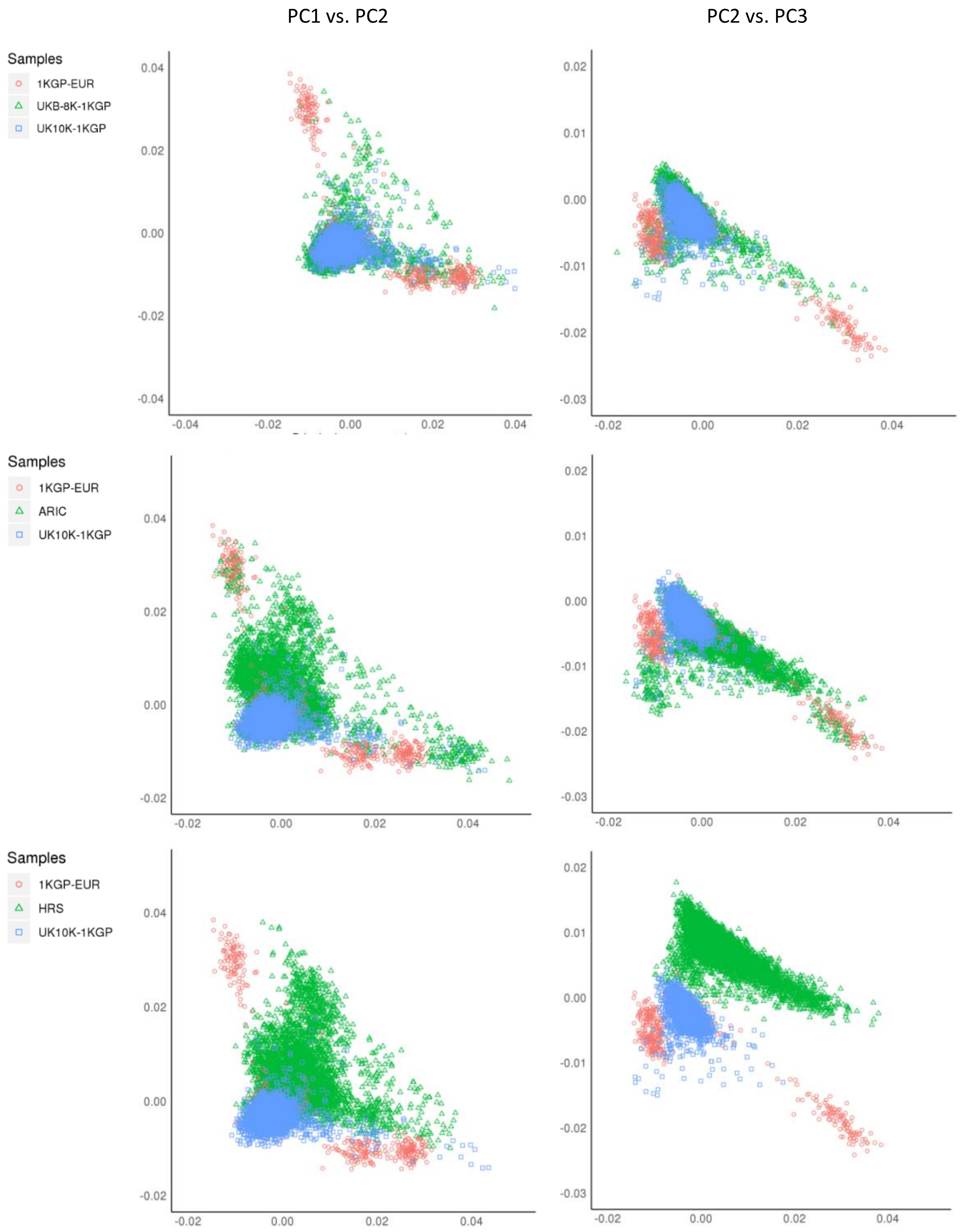


**Supplementary Figure 6. Principal component analysis of the LD reference samples.** We combined the genotype data of HRS, ARIC and UKB-8K-1KGP with those of 1KGP-EUR and performed a principal component analysis (PCA) using the variants with MAF>0.05. The panels on the left-hand side show the plots of PC1 against PC2, and those on the right-hand side show the plots of PC3 against PC2. As expected, UKB-8K-1KGP shows a greater overlap with UK10K-1KGP than ARC or HRS.

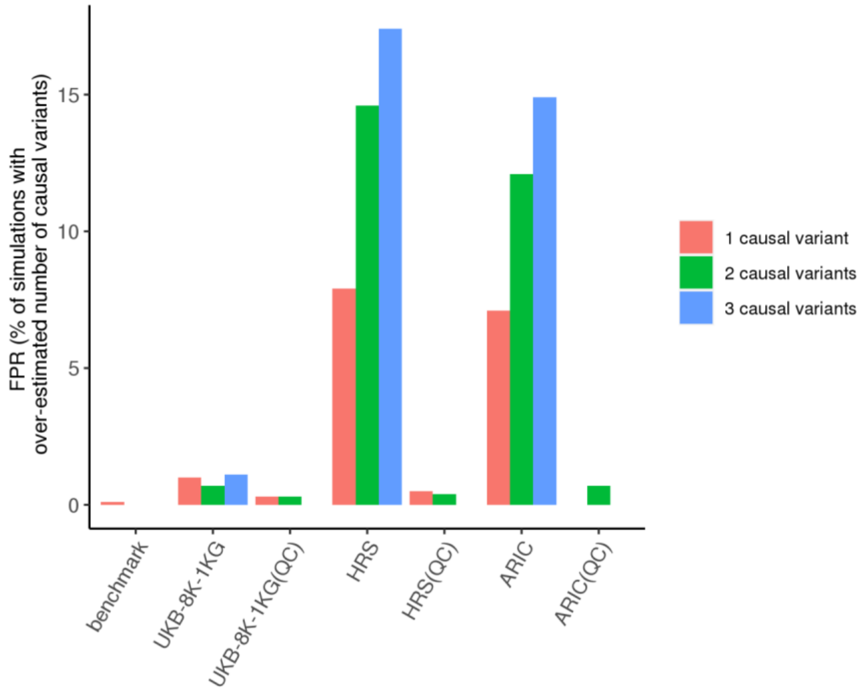


**Supplementary Figure 7. FPR of COJO with and without DENTIST QC in simulations with one, two or three causal variant(s) at one locus.** Based on simulations with 1, 2 or 3 causal variant(s), we assessed the FPRs of COJO when performed with and without DENTIST-based QC (FPR is defined as the proportion of simulations in which the number of COJO signals is larger than the number of causal variants). The x-axis labels indicate the LD reference samples, and those performed after DENTIST QC are labeled with "QC" in the parentheses.

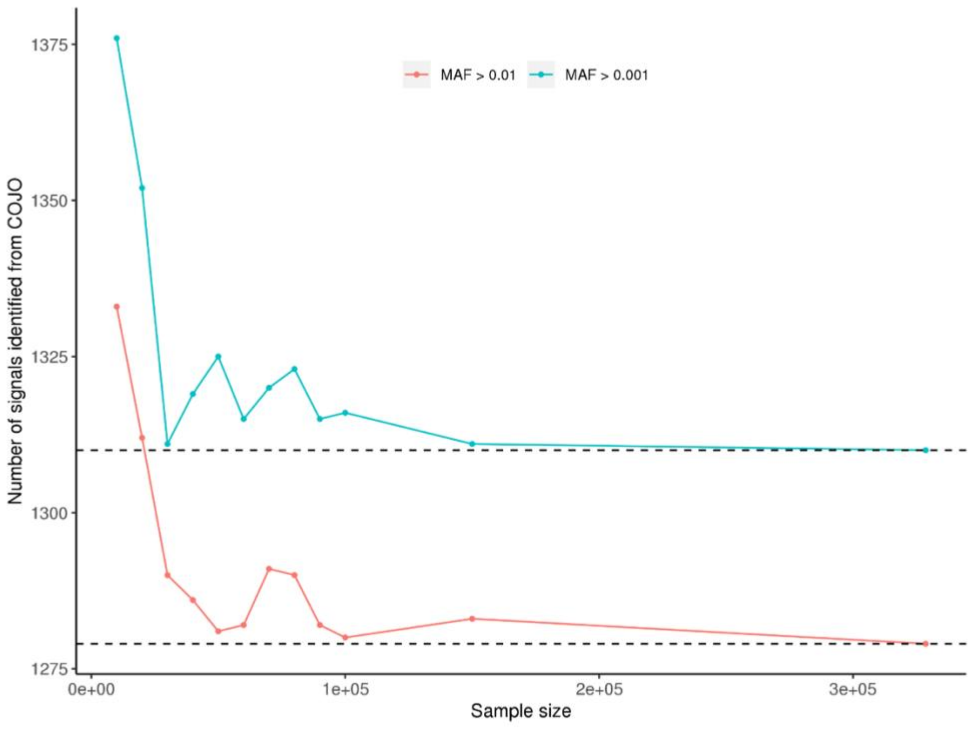


**Supplementary Figure 8. The change of the number COJO signals with the increase of the in-sample LD reference sample size.** By subsetting the discovery GWAS sample (328,577 unrelated UKB participants), we created 11 in-sample LD references with sample sizes ranging from 10,000 to 150,000. As the reference sample size ($n_{ref}$) increases, the number of signals identified from COJO with MAF>0.01 first decreases sharply from 1,333 at $n_{ref}$ =10,000 to 1,290 at $n_{ref}$ =30,000, before stabilizing in a range between 1,280 and 1,291, close to the benchmark of 1,279 signals identified using the whole discovery GWAS sample as the reference (the right-most dot on the plot). The COJO results for variants with MAF > 0.001 show a similar pattern, i.e., the number of COJO signals deceases sharply when $n_{ref}$ increases from 10,000 to 30,000 and stabilizes in the range between 1,310 and 1,325.

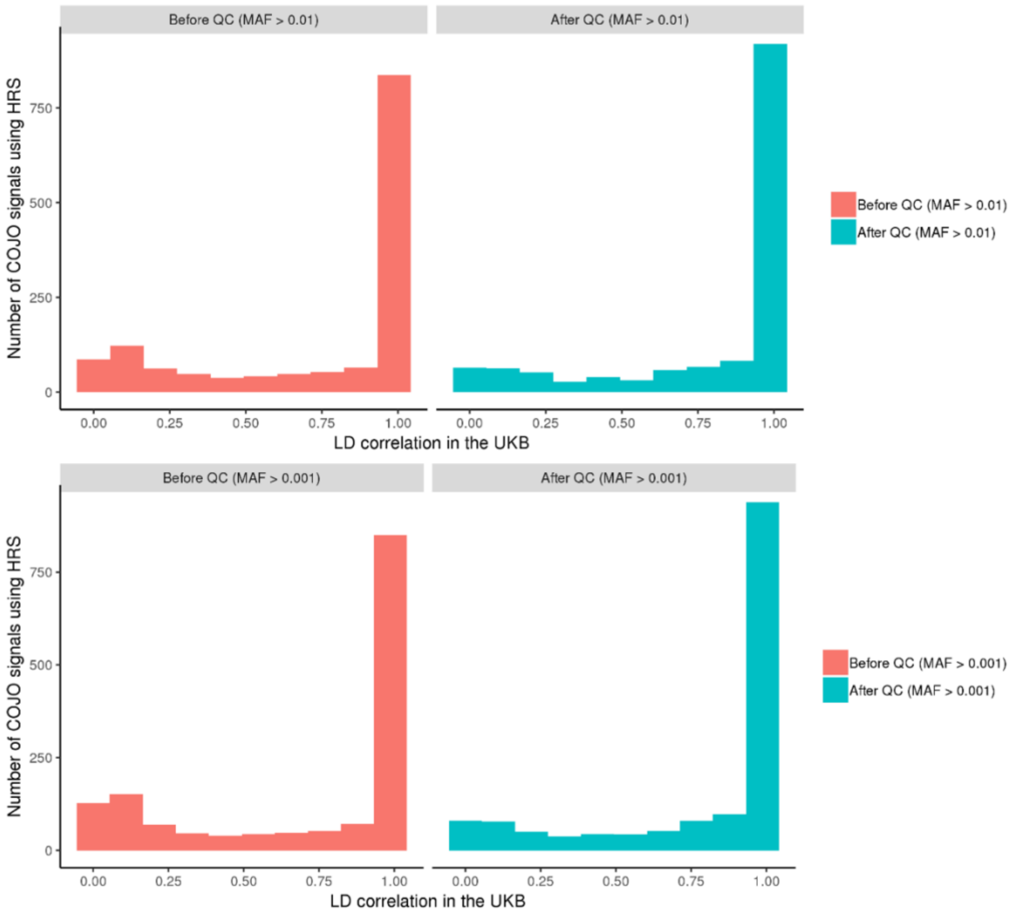


**Supplementary Figure 9. LD correlations between the COJO signals identified using HRS as the reference and their best matching signals identified from the benchmark analysis.** Shown are the LD correlations between the COJO signals for height, identified using HRS as the LD reference before and after DENTIST-based QC, and their best matching signals identified from the benchmark analysis using the whole discovery GWAS (UKBv3-329K) as the LD reference. For variants with MAF >0.01, a few signals identified using HRS shown on the top-right panel show weak LD ($r^2 < 0.3$) with the benchmark signals. After DENTIST-QC, the number of poorly correlated signals is greatly reduced as shown on the top-right panel ($r^2 < 0.3$). On the other hand, the DENTIST-based QC leads to more strong correlations ($r^2 > 0.9$). A similar pattern is observed for variants with MAF>0.001 (shown in the two bottom panels).

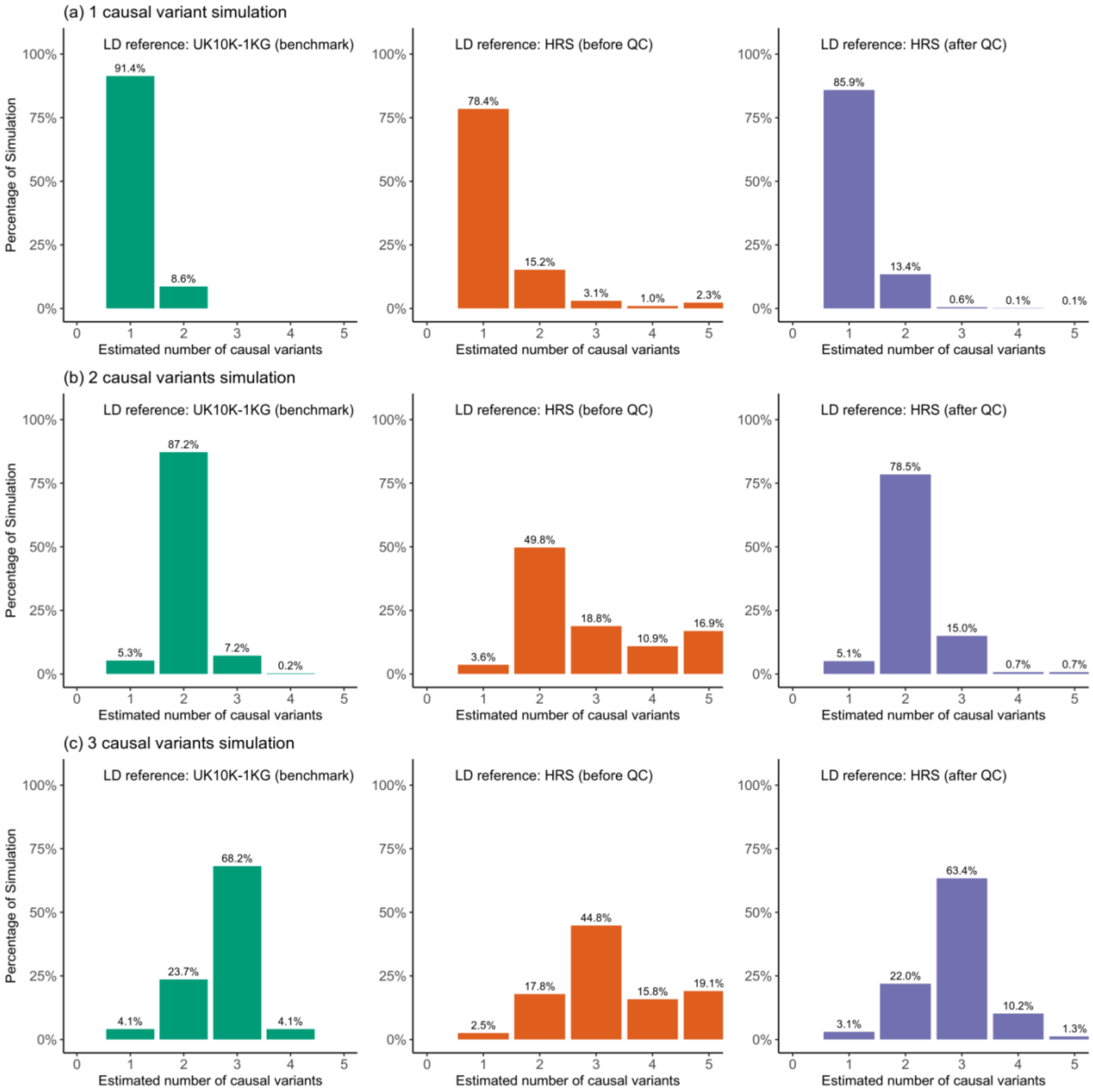


**Supplementary Figure 10**. **FINEMAP analysis using out-of-sample LD from HRS with or without DENTIST QC compared to the benchmark.** We assessed the performance of FINEMAP by comparing the estimated number of causal variants to the true number of causal variants in simulations with 1, 2 or 3 causal variants at on one locus. Each scenario was repeated 200 times with the causal variants resampled in each replicate. Plotted is the proportion of simulation replicates in which the estimated number of causal variants from FINEMAP was consistent with the true number. Benchmark: FINEMAP analysis using in-sample LD (without DENTIST QC).

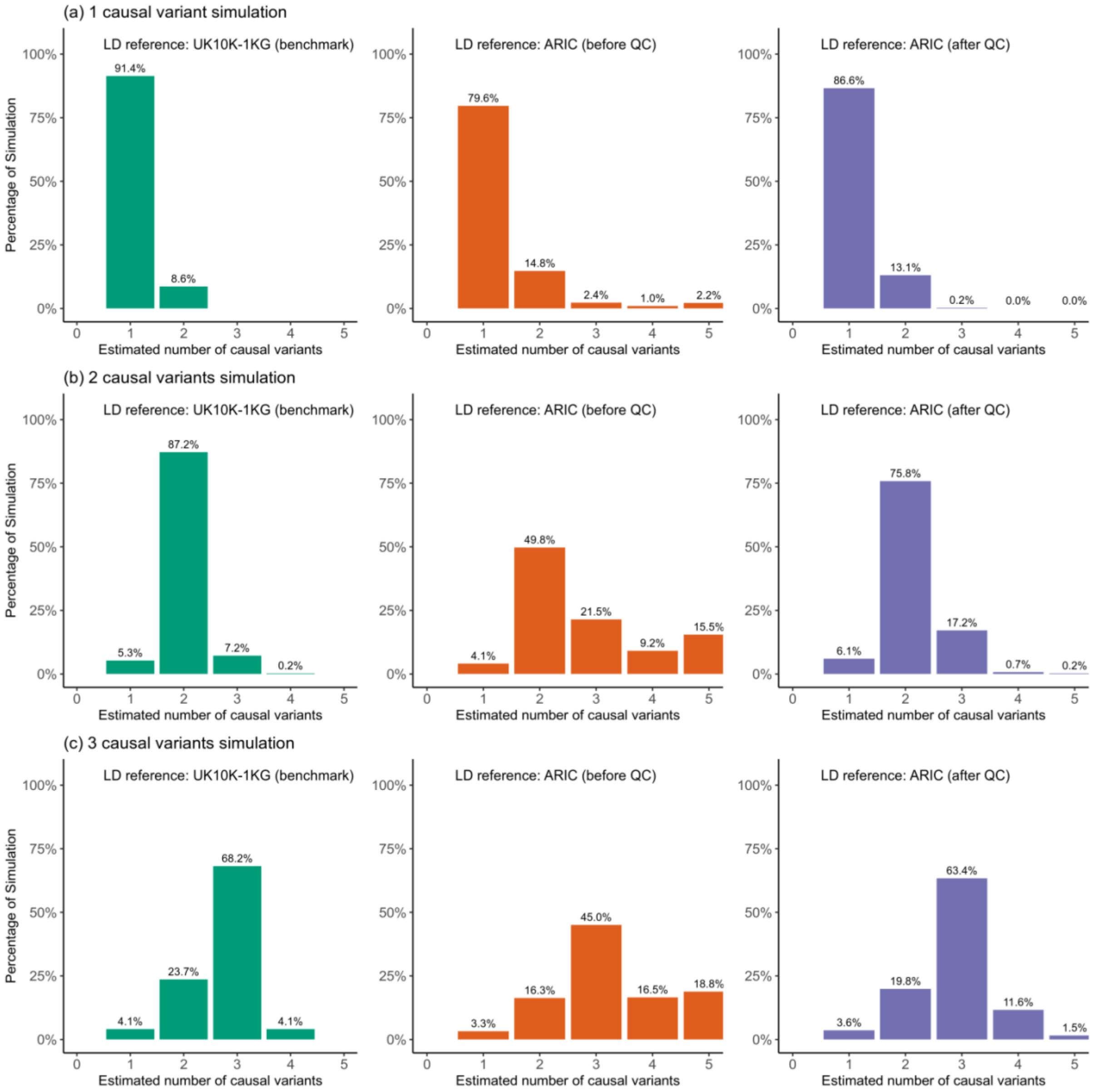


**Supplementary Figure 11**. **FINEMAP analysis using out-of-sample LD from ARIC with or without DENTIST QC compared to the benchmark.** We assessed the performance of FINEMAP by comparing the estimated number of causal variants to the true number of causal variants in simulations with 1, 2 or 3 causal variants at on one locus. Each scenario was repeated 200 times with the causal variants resampled in each replicate. Plotted is the proportion of simulation replicates in which the estimated number of causal variants from FINEMAP was consistent with the true number. Benchmark: FINEMAP analysis using in-sample LD (without DENTIST QC).

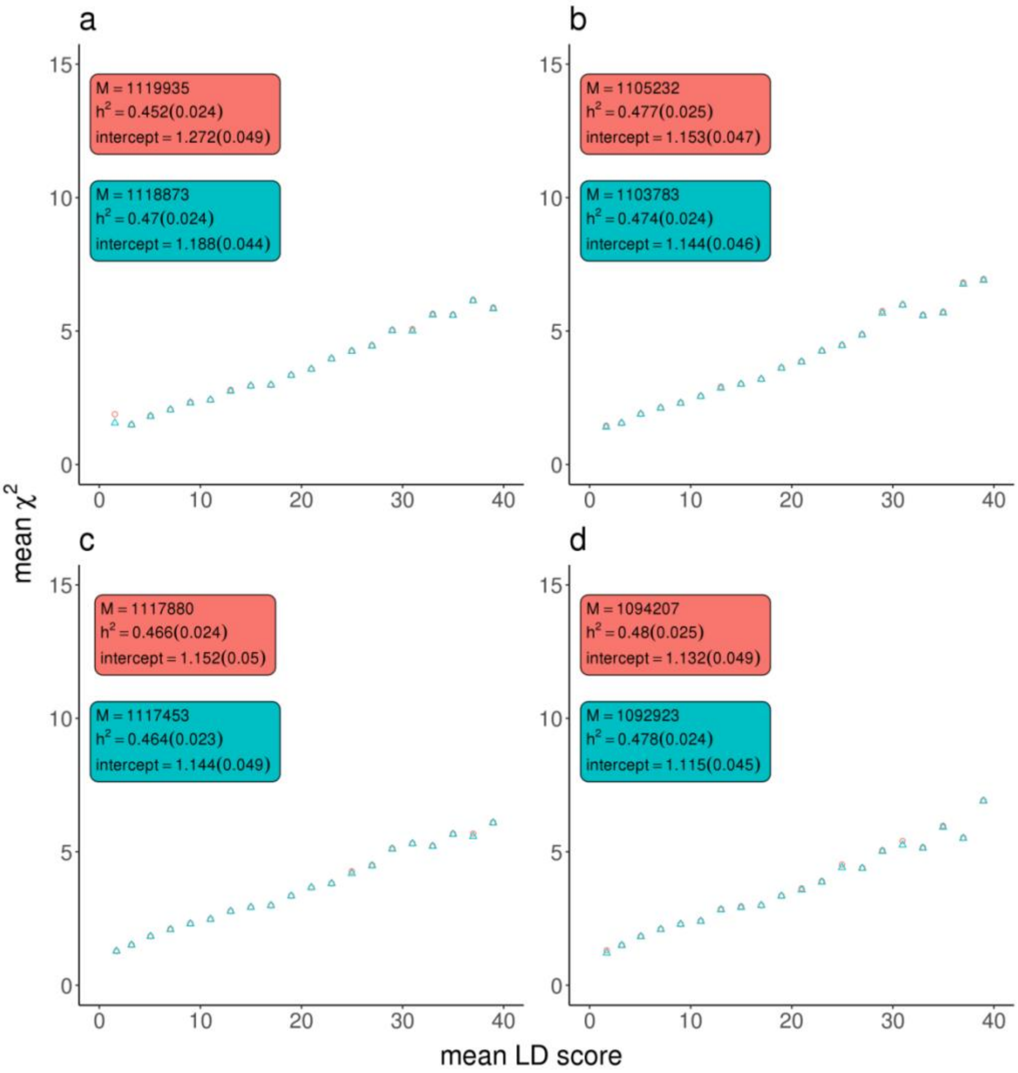


**Supplementary Figure 12. The effect of DENTIST-based QC on LDSC analysis of the UKB height summary data.** We assessed the effect of DENTIST on LDSC when different LD references were used, including **a**) HRS, **b**) ARIC, **c**) UKB-8K-1KGP, and **d**) UK10K-WGS. For each reference sample, LDSC was performed before and after DENTIST-based QC, and the corresponding results are shown in the red and cyan text boxes, respectively, on each plot. The variants are binned by their LD scores. Each dot on the plots represents the mean LD score value of each bin on the x-axis and the mean $\chi^2$ value on the y-axis, with those before and after DENTIST-based QC in red and cyan colors respectively. In the textbox, "M" represents the number of variants, "$h_{SNP}^2$" represents the estimate of SNP-based heritability, and "intercept" represents the LDSC intercept, with the corresponding standard errors given the parentheses.

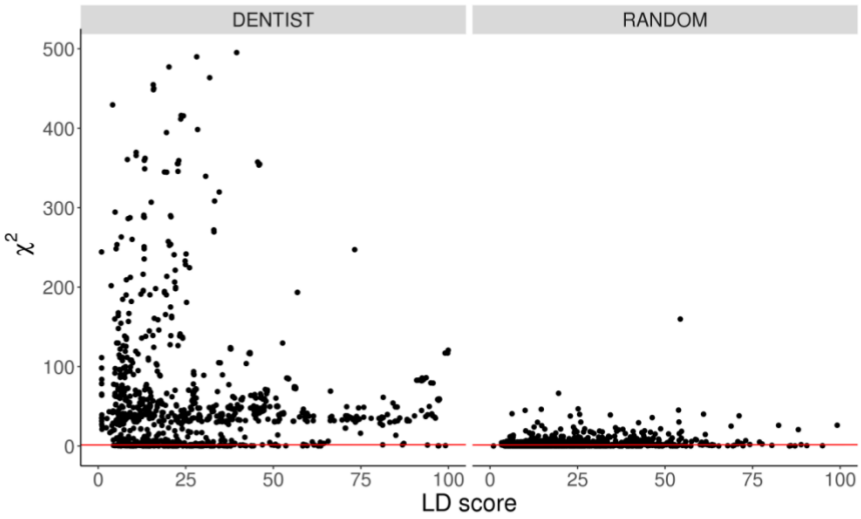


**Supplementary Figure 13. A plot of LD scores of the variants detected by DENTIST against their GWAS test-statistics for height in the UKB.** This plot shows the LD scores and $\chi^2$ statistics of variants identified from a DENTIST analysis of the UKB height GWAS data (n = 328,577 unrelated UKB participants) using HRS as the LD reference compared to variants selected at random. The red line represents the regression line with its intercept and slope estimated from the LDSC analysis of all the variants.

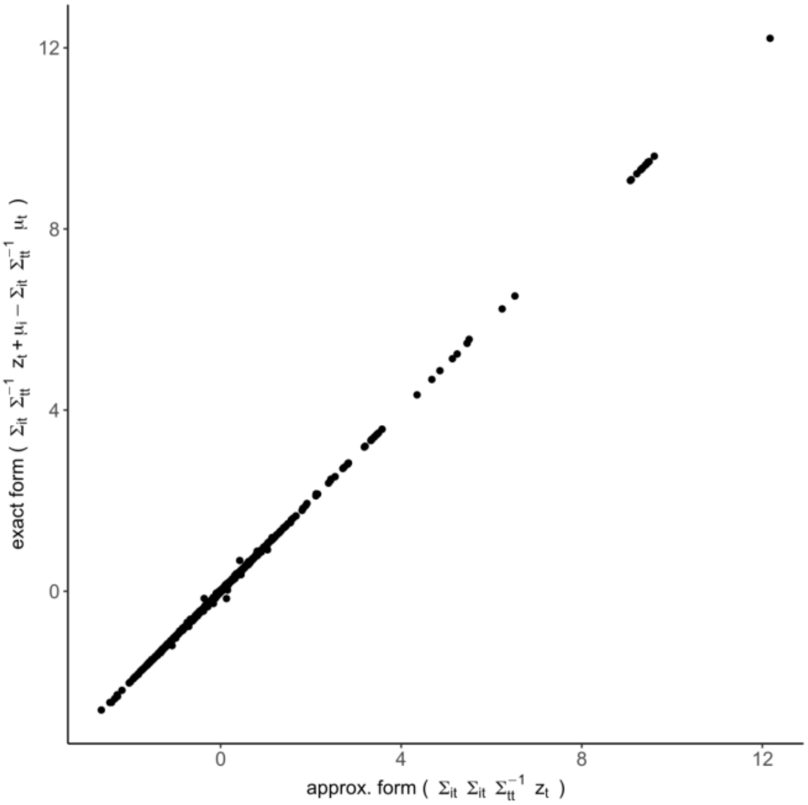


**Supplementary Figure 14. Comparison between imputed z-score derived under null model (approx. form) and imputation under fixed-effect model (exact form).** We used UK10K.WGS sample to simulate a phenotype caused by a variant *c* with effect denoted by $\lambda_c$ and heritibility fixed to $h^2 = 0.05$. Therefore, the expected values for each marginal effect measured in z-score, $\mu_i$, equals to $\Sigma_{ic}\lambda_c$, where $\Sigma_{ic}$ is the LD correlation between variants *i* and *c*.

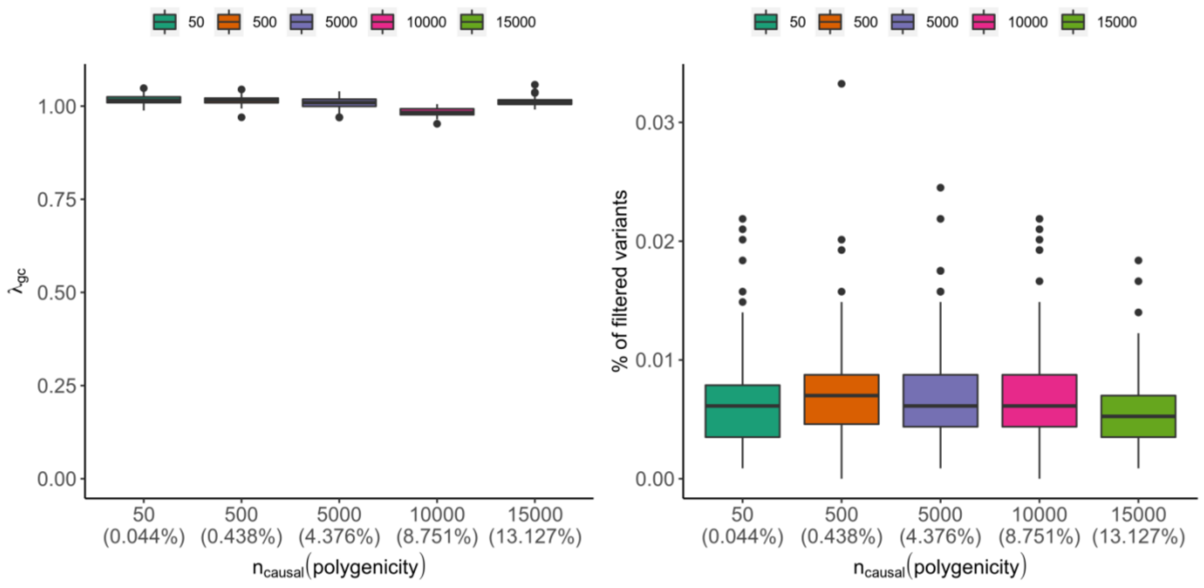


**Supplementary Figure 15. The impact of the genetic architecture of the trait on DENTIST.** We assessed DENTIST under a range of MAF- and LD-dependent genetic architectures by simulation using data on chromosome 22 from UK10K-WGS, ranging from oligogenic ($m_{\text{causal}}$ = 50) to very polygenic ones ($m_{\text{causal}}$ =15,000, accounting for 13.1% of all the variants). The effects were sampled from $N(0, [2p(1-p)]^{-1})$ with $p$ being the MAF, and the trait heritability $h^2$ was set to 20%. The causal variants were sampled from a set of variants with larger LD scores than average, to generate a LD-dependent genetic architecture. Each scenario was repeated 100 times, with the causal variants and their effect sizes resampled in each replicate. No errors were simulated in this scenario. Shown in panels a) and b) are the inflation factor of the DENTIST test statistic and the proportion of variants removed by DENTIST, respectively. This is no relationship between the inflation factor of DENTIST test statistic (or the proportion of variants filtered by DENTIST) and the number of causal variants simulated, and all these results are similar to those from simulations based on a simple genetic architecture as presented in Supplementary Figure 1, suggesting the robustness of DENTIST to the genetic architecture of the trait.

### Supplementary Tables

**Supplementary Table 1.** A summary of the GWAS data sets used in the present study.

| | Genotyping platform | Imputation reference | HWE P-value threshold | MAC/MAF threshold | Number of variants with MAF > 0.01 | Number of variants with MAF > 0.001 |
|---|---|---|---|---|---|---|
| HRS (n = 8,557) | Illumina Omni 2.5 | KG3 | $10^{-6}$ | MAC of 5 | 11,326,629 | 15,608,163 |
| ARIC (n=7703) | Affymetrix 6.0 | KG3 | $10^{-6}$ | MAC of 5 | 8,627,204 | 13,356,572 |
| UK10K-WGS (n=3,642) | WGS (> 30x coverage) | NA | $10^{-6}$ | MAC of 3 | 8,325,293 | 12,798,760 |
| UK10K-1KGP (n=3642) | Illumina Core Exome array | KG3 | $10^{-6}$ | MAC of 3 | 9,351,188 | 14,398,774 |
| 1KG-EUR (n= 503) | WGS (~7.4x coverage) | NA | $10^{-6}$ | MAF of 0.01 | 9,417,956 | 9,417,956 |
| HRS-3K (n=3,000) at chr22 | Illumina Omni 2.5 | KG3 | $10^{-6}$ | MAF of 0.01 | 45,901 | 45,901 |
| UKBv3 (n=348577) | Affymetrix Axiom | HRC UK10K | $10^{-6}$ | MAF of 0.0001 | 8,569,717 | 13,072,891 |
| UKBv3-329K at chr22 (n=34,8580) | Affymetrix Axiom | HRC UK10K | $10^{-6}$ | MAF of 0.0001 | 8,569,717 | 13,072,891 |
| UKBv3-8K (n=8000) | Affymetrix Axiom | HRC UK10K | $10^{-6}$ | MAF of 0.0001 | 9,351,188 | 14,398,774 |
| UKBv3-20K (n=20,000) | Affymetrix Axiom | HRC UK10K | $10^{-6}$ | MAF of 0.0001 | 9,351,188 | 14,398,774 |
| UKB-8K-1KGP (n=8,000) | Affymetrix Axiom | KG3 | $10^{-6}$ | MAF of 0.0001 | 7,349,687 | 9,972,916 |

1KGP: 1000 Genome Project Phase III data. WGS: whole genome sequencing. HRC: Haplotype Reference Consortium. HM3: HapMap 3 Project.

**Supplementary Table 2.** Proportion of simulated erroneous variants identified by standard QC.

| LD reference | %variants with allelic errors detected by the ΔAF test | | | %variants with genotyping errors detected by the HWE test |
|---|---|---|---|---|
| | MAF ≥ 0.01 | MAF > 0.4 | MAF > 0.45 | |
| UKB-8K-1KGP | 95±1.0 | 57±6.8 | 16±6.9 | 45±2.1 |
| UK10K–WGS | 94±1.1 | 50±6.6 | / | 52±2.3 |

Shown are mean ± standard error.

**Supplementary Table 3.** Power in detecting simulated genotyping and allelic errors in UK10K-WGS using out-of-sample LD from UKB-8K-1KGP.

| | **Variants stratified into bins by \|z-score\| or p-value** | | | | | | **All** |
|---|---|---|---|---|---|---|---|
| | \|z-score\| [0, 1) | \|z-score\| [1, 2) | \|z-score\| [2, 3) | \|z-score\| [3, 4) | \|z-score\| >4 | *P*-value $<5\times10^{-8}$ | |
| Allelic error | 57.4±1.2 | 99.2±0.3 | 100±0.0 | 100±0.0 | 100±0.0 | 100±0.0 | **73.4±0.9** |
| 5% genotyping error rate | 18.4±0.5 | 21.8±0.7 | 30.7±1.7 | 49.2±4.4 | 56.9±5.8 | 84.8±6.2 | 20.9±0.4 |
| 10% genotyping error rate | 26.2±0.6 | 31.4±1.0 | 47.6±2.2 | 61.3±5.4 | 92.3±3.7 | 100±0.0 | 29.9±0.5 |
| 15% genotyping error rate | 29.8±0.7 | 38.7±1.2 | 52.1±2.6 | 84.7±3.9 | 92.3±5.2 | 100±0.0 | 34.8±0.6 |
| All genotyping errors | 23.7±0.3 | 29.0±0.5 | 40.9±1.2 | 62.8±2.8 | 75.3±3.5 | 93.5±2.8 | **27.2±0.3** |

"All genotyping errors" refers to all the genotyping errors regardless of genotyping error rate setting. Shown are mean ± standard error.

**Supplementary Table 4.** Proportion of variants removed by different QC strategies in the simulations based on the UK10K-WGS data.

| Reference sample | Standard QC | DENTIST-based QC |
|---|---|---|
| **Among all the variants** | | |
| UKB-8K-1KGP | 0.6 ± 0.01% | 0.20±0.02% |
| UK10K-WGS | 0.7 ± 0.01% | 0.13±0.01% |
| **Among the causal variants** | | |
| UKB-8K-1KGP | 0.44 ± 1.1% | 0.99±1.4% |
| UK10K-WGS | 0.44 ± 1.1% | 0.19±0.63% |

**Supplementary Table 5.** Fold enrichment in probability of detecting an erroneous variant using DENTIST before and after standard QCs.

| | **Allelic errors** | **Genotyping errors** | | | |
|---|---|---|---|---|---|
| | | Genotyping error rate = 5% | Genotyping error rate = 15% | Genotyping error rate = 15% | All |
| **Using UKB-8K-1KGP as the reference (out-of-sample LD)** | | | | | |
| DENTIST before standard QCs | 82 ± 4.7 | 33 ± 4.9 | 44 ± 6.3 | 49 ± 5.8 | 42 ± 3.9 |
| DENTIST after standard QCs | 366±58.7 | 105±21.8 | 150±26.4 | 176±34.4 | 137±17.0 |
| **Using UK10K-WGS as the reference (in-sample LD)** | | | | | |
| DENTIST before standard QCs | 103 ± 4.0 | 51 ± 5.0 | 64 ± 5.9 | 69 ± 5.4 | 61 ± 3.0 |
| DENTIST after standard QCs | 575±87.8 | 325±51.9 | 212±32.8 | 286±40.2 | 261±27.7 |

"All" refers to all the genotyping errors regardless of the genotyping error rate setting. Shown are mean ± standard error.

**Supplementary Table 6.** Power of DENTIST in detecting simulated genotyping and allelic errors using the UK10K-WGS sample as the LD reference.

| | **\|z-score\| [0, 1)** | **\|z-score\| [1, 2)** | **\|z-score\| [2, 3)** | **\|z-score\| [3, 4)** | **\|z-score\| >4** | **\|z-score\| >5.45** | **\|z-score\| >0** |
|---|---|---|---|---|---|---|---|
| **Allelic errors** | 59.8±1.1 | 98.3±0.4 | 100.0±0.0 | 100.0±0.0 | 100.0±0.0 | 100.0±0.0 | 74.6±0.8 |
| **Genotyping errors (5%)** | 24.8±0.5 | 29.7±0.7 | 35.7±1.6 | 53.5±4.0 | 64.3±5.2 | 81.8±5.8 | 27.6±0.4 |
| **Genotyping errors (10%)** | 33.9±0.6 | 39.6±0.9 | 53.2±2.0 | 77.9±4.1 | 96.7±2.3 | 97.0±3.0 | 37.7±0.5 |
| **Genotyping errors (15%)** | 37.2±0.7 | 45.3±1.1 | 58.9±2.4 | 79.4±4.1 | 96.9±3.1 | 100.0±0.0 | 41.7±0.6 |
| **All genotyping errors** | 30.6±0.3 | 36.6±0.5 | 46.3±1.1 | 67.6±2.5 | 81.3±2.9 | 91.0±2.9 | 34.1±0.3 |

"All genotyping errors" refer to all the genotyping errors regardless of the genotyping error rate setting. |z-score| >5.45 corresponds to P-value $<5\times10^{-8}$. Shown are mean ± standard error.

**Supplementary Table 7.** The loss of power in GWAS due to imperfect tagging of the causal variants.

| | **Common** | **Rare** |
|---|---|---|
| **Variant tagging (absolute value of Pearson's correlation between imputed and sequenced genotypes)** | 0.77± 0.007 | 0.46±0.013 |
| **Proportion of simulations containing ≥1 genome-wide significant signals** | 89% | 50% |

Shown are mean ± standard error.

**Supplementary Table 8.** COJO analysis of the UKB GWAS summary data for height using the whole discovery or a subset of the discovery sample as the LD reference.

| Sample size | MAF > 0.01 | MAF > 0.001 |
|---|---|---|
| 10000 | 1,333 | 1,376 |
| 20000 | 1,312 | 1,352 |
| 30000 | 1,290 | 1,311 |
| 40000 | 1,286 | 1,319 |
| 50000 | 1,281 | 1,325 |
| 60000 | 1,282 | 1,315 |
| 70000 | 1,291 | 1,320 |
| 80000 | 1,290 | 1,322 |
| 90000 | 1,282 | 1,315 |
| 100000 | 1,280 | 1,316 |
| 150000 | 1,283 | 1,311 |
| All (n=328577) | 1,279 | 1,310 |

**Supplementary Table 9.** The accuracy ($r^2$) of using the COJO signals identified from the UKB to predict height in HRS.

| | LD reference for COJO and DENTIST | | |
|---|---|---|---|
| | **ARIC** | **UKBv3-8K** | **UKBv3-20K** |
| Using COJO signals at MAF > 0.01 | | | |
| COJO only | 0.18±0.007 | 0.19±0.007 | 0.19±0.007 |
| COJO + DENTIST | 0.19±0.007 | 0.19±0.007 | 0.19±0.007 |
| Using COJO signals at MAF > 0.001 | | | |
| COJO only | 0.18±0.007 | 0.19±0.007 | 0.19±0.008 |
| COJO + DENTIST | 0.19±0.007 | 0.19±0.007 | 0.19±0.007 |

Note: we constructed a polygenic score for height in HRS based on the COJO signals identified from the UKB height data using three different references. The numbers of COJO signals identified using different LD references before and after DENTIST can be found in Table 2. Shown in this table are mean ± standard error of the correlation squared between the PRS and the real height phenotype in the HRS accounting for the effects of age, sex and the first 10 PCs.

**Supplementary Table 10.** Proportion of variants in the UKB height GWAS data removed by DENTIST using out-of-sample references.

| | **MAF > 0.01** | **MAF > 0.001** |
|---|---|---|
| **UKBv8k** | 0.04% | 0.04% |
| **UKBv20k** | 0.03% | 0.11% |
| **ARIC** | 0.33% | 1.4% |
| **HRS** | 0.34% | 1.3% |
| **UK10K-WGS** | 0.16% | 0.73% |

**Supplementary Table 11.** A summary of the 10 published GWAS summary data sets used in this study.

| Trait (abbreviation) | $n$ | $n_{\text{cases}}$ | $n_{\text{controls}}$ | Year | PUBMED ID |
|---|---|---|---|---|---|
| Educational Attainment (EA) | 766,345 | / | / | 2018 | 30038396 |
| Coronary Artery Disease (CAD) | 547,261 | 122,733 | 424,528 | 2017 | 29212778 |
| Type 2 Diabetes(T2D) | 898,130 | 74,124 | 824,006 | 2018 | 30297969 |
| Crohn's Disease (CD) | 20,883 | 5,956 | 14,927 | 2015 | 26192919 |
| Major Depressive Disorder (MDD) | 480,359 | 135,458 | 344,901 | 2018 | 29700475 |
| Schizophrenia (SCZ) | 35,802 | 11,260 | 24,542 | 2018 | 29483656 |
| Ovarian Cancer (OC) | 66,450 | 25,509 | 40,941 | 2018 | 28346442 |
| Breast Cancer (BC) | 228,951 | 12,2977 | 105,974 | 2018 | 29059683 |
| Height | ~700,000 | / | / | 2018 | 30124842 |
| Body Mass Index (BMI) | ~700,000 | / | / | 2018 | 30124842 |

$n$: sample size; $n_{\text{cases}}$ and $n_{\text{controls}}$: numbers of cases and controls, respectively.

**Supplementary Table 12.** Proportion of variants removed in DENTIST analyses of the 10 published GWAS summary data sets using different LD references.

| Trait | LD reference | | |
|---|---|---|---|
| | **ARIC** | **HRS** | **UKBv3.8K** |
| BMI | **0.07%** | **0.05%** | **0.05%** |
| Height | 0.67% | 0.62% | 0.75% |
| SCZ | 0.13% | 0.17% | 0.63% |
| EA | 0.21% | 0.23% | 0.22% |
| CAD | **0.86%** | **0.92%** | **1.46%** |
| T2D | 0.23% | 0.24% | 0.22% |
| CD | 0.30% | 0.33% | 0.66% |
| MDD | 0.57% | 0.59% | 0.72% |
| BC | 0.15% | 0.17% | 0.21% |
| OC | 0.09% | 0.10% | 0.09% |

**Supplementary Table 13.** Number of COJO signals from the COJO analysis of the 10 published GWAS data sets with and without DENTIST-based QC.

| Trait | DENTIST QC | LD reference for DENTIST and COJO | | |
|---|---|---|---|---|
| | | **ARIC** | **HRS** | **UKBv3.8K** |
| BMI | No | 728 | 744 | 727 |
| | Yes | 726 (-0.3%) | 734 (-1.3%) | 731 (+0.6%) |
| Height | No | 3,145 | 3,109 | 3,040 |
| | Yes | 3,088 (-1.8%) | 3,046 (-2.0%) | 2,990 (-1.6%) |
| SCZ | No | 155 | 150 | 142 |
| | Yes | 148 (-4.5%) | 145 (-3.3%) | 141 (-0.7%) |
| EA | No | 480 | 485 | 467 |
| | Yes | 480 (-0.0%) | 479 (-1.2%) | 466 (-0.2%) |
| CAD | No | 192 | 199 | 189 |
| | Yes | **168 (-12.5%)** | **178 (-10.6%)** | **170 (-10.1%)** |
| T2D | No | 274 | 263 | 267 |
| | Yes | 268 (-2.2%) | 262 (-0.4%) | 268 (0.4%) |
| CD | No | 64 | 66 | 63 |
| | Yes | 61 (-4.7%) | 63 (-4.5%) | 62 (-1.6%) |
| MDD | No | 45 | 45 | 44 |
| | Yes | **45 (0%)** | **45 (0%)** | **42 (-4.5%)** |
| BC | No | 194 | 193 | 181 |
| | Yes | 182 (-6.2%) | 181 (-6.2%) | 175 (-3.3%) |
| OC | No | 12 | 12 | 12 |
| | Yes | **12 (0%)** | **12 (0%)** | **12 (0%)** |

The numbers in parentheses are the percentages of decrease (-) or increase (+) in the number of COJO signals after DENTIST-QC.

**Supplementary Table 14.** Estimates from LDSC analyses of the UKB height GWAS summary data with and without DENTIST-based QC.

| | Number of variants | One-step approach | | Two-step approach | |
|---|---|---|---|---|---|
| | | $h^2_{SNP}$ | Intercept | $h^2_{SNP}$ | Intercept |
| **Reference = the discovery GWAS sample** | | | | | |
| **Benchmark** | 1,114,780 | 0.46 (0.023) | 1.13 (0.049) | 0.41(0.017) | 1.34 (0.030) |
| **Reference = HRS** | | | | | |
| **Without DENTIST** | 1,118,873 | 0.45 (0.023) | 1.27 (0.049) | 0.42 (0.018) | 1.37 (0.029) |
| **With DENTIST** | 1,116,249 | 0.47 (0.023) | 1.18 (0.043) | 0.41 (0.017) | 1.37 (0.029) |
| **Reference = ARIC** | | | | | |
| **Without DENTIST** | 1,105,232 | 0.47 (0.025) | 1.15 (0.047) | 0.42 (0.018) | 1.34 (0.030) |
| **With DENTIST** | 1,103,783 | 0.47 (0.024) | 1.14 (0.047) | 0.41 (0.018) | 1.34 (0.030) |
| **Reference = UKB-8K-1KGP** | | | | | |
| **Without DENTIST** | 1,117,880 | 0.46(0.024) | 1.15(0.049) | 0.41 (0.018) | 1.34 (0.031) |
| **With DENTIST** | 1,117,453 | 0.46(0.023) | 1.14(0.048) | 0.40 (0.017) | 1.34 (0.031) |
| **Reference = UK10K-WGS** | | | | | |
| **Without DENTIST** | 1,074,399 | 0.47 (0.024) | 1.13(0.049) | 0.42 (0.018) | 1.33 (0.030) |
| **With DENTIST** | 1,092,923 | 0.47 (0.023) | 1.11 (0.044) | 0.41 (0.017) | 1.33(0.030) |
| **Reference = 1KGP-EUR** | | | | | |
| **Without DENTIST** | 1,133,151 | 0.48 (0.024) | 1.15 (0.047) | 0.43 (0.018) | 1.33 (0.028) |
| **With DENTIST** | 1,124,951 | 0.471 (0.023) | 1.15 (0.048) | 0.42 (0.018) | 1.32 (0.023) |

Standard errors are given in the parentheses.

**Supplementary Table 15.** LDSC results for the 10 published GWAS summary data sets with or without DENTIST-based QC.

| Trait | DENTIST QC | LD reference for DENTIST and LDSC | | |
|---|---|---|---|---|
| | | ARIC | UK10K-WGS | UKBv3-8K |
| BMI | No | $m$=969,753;<br>$h_{SNP}^2$=0.172(0.006)<br>intercept=1.034(0.026) | $m$=955,844;<br>$h_{SNP}^2$=0.173(0.006)<br>intercept=1.008(0.025) | $m$=976,507;<br>$h_{SNP}^2$= 0.168 (0.006)<br>intercept= 1.026 (0.024) |
| | Yes | $m$=968,979;<br>$h_{SNP}^2$=0.170(0.005)<br>intercept=1.034(0.024) | $m$=955,229;<br>$h_{SNP}^2$=0.172(0.006)<br>intercept=1.009(0.025) | $m$=975,871;<br>$h_{SNP}^2$=0.164 (0.005)<br>intercept= 1.030 (0.024) |
| Height | No | $m$=965,367;<br>$h_{SNP}^2$=0.511(0.026)<br>intercept=1.368(0.124) | $m$=951,213;<br>$h_{SNP}^2$=0.510(0.026)<br>intercept=1.306(0.123) | $m$=971,821;<br>$h_{SNP}^2$=0.496(0.025)<br>intercept=1.347(0.122) |
| | Yes | $m$=958,942;<br>$h_{SNP}^2$=0.472(0.024)<br>intercept=1.275(0.008) | $m$=943,562;<br>$h_{SNP}^2$=0.46(0.021)<br>intercept=1.205(0.080) | $m$=964,237;<br>$h_{SNP}^2$=0.447(0.020)<br>intercept=1.234(0.076) |
| SCZ | No | $m$=1,121,978;<br>$h_{SNP}^2$=0.658(0.027)<br>intercept=1.036(0.013) | $m$=1,107,162;<br>$h_{SNP}^2$=0.651(0.025)<br>intercept=1.032(0.013) | $m$=1,092,944;<br>$h_{SNP}^2$=0.632 (0.025)<br>intercept= 1.038 (0.014) |
| | Yes | $m$=1,120,942;<br>$h_{SNP}^2$=0.656(0.027)<br>intercept=1.036(0.013) | $m$=1,099,585;<br>$h_{SNP}^2$=0.646(0.023)<br>intercept=1.032(0.013) | $m$=1,089,005;<br>$h_{SNP}^2$=0.628 (0.023)<br>intercept=1.039 (0.013) |
| EA | No | $m$=1138090;<br>$h_{SNP}^2$=0.113(0.003)<br>intercept=0.961(0.016) | $m$=1,110,625;<br>$h_{SNP}^2$=0.114(0.003)<br>intercept=0.935(0.016) | $m$=1,097,272;<br>$h_{SNP}^2$= 0.110 (0.003)<br>intercept= 0.948 (0.017) |
| | Yes | $m$=1,136,715;<br>$h_{SNP}^2$=0.113(0.003)<br>intercept=0.960(0.016) | $m$=1,106,705;<br>$h_{SNP}^2$=0.113(0.003)<br>intercept=0.932(0.016) | $m$=1,095,657;<br>$h_{SNP}^2$=0.11 (0.003)<br>intercept= 0.949(0.017) |
| CAD | No | $m$=1,147,517<br>$h_{SNP}^2$=0.060(0.004)<br>intercept=0.975(0.017) | $m$=1,131,860<br>$h_{SNP}^2$=0.060(0.004)<br>intercept=0.971(0.015) | $m$=1,115,519;<br>$h_{SNP}^2$=0.053 (0.004)<br>intercept=0.975 (0.016) |
| | Yes | $m$=1,124,598;<br>$h_{SNP}^2$=0.054(0.003)<br>intercept=0.977(0.016) | $m$=1,091,284;<br>$h_{SNP}^2$=0.054(0.003)<br>intercept=0.967(0.013) | $m$=1,085,336<br>$h_{SNP}^2$=0.053 (0.003)<br>intercept=0.976 (0.014) |
| T2D | No | $m$=1,103,665;<br>$h_{SNP}^2$=0.193(0.009)<br>intercept=1.077(0.028) | $m$=1,090,573;<br>$h_{SNP}^2$=0.192(0.009)<br>intercept=1.071(0.028) | $m$=1,115,665;<br>$h_{SNP}^2$=0.186(0.009)<br>intercept=1.080(0.029) |
| | Yes | $m$=1,101,989;<br>$h_{SNP}^2$=0.191(0.009)<br>intercept=1.070(0.025) | $m$=1,087,387;<br>$h_{SNP}^2$=0.190(0.009)<br>intercept=1.065(0.026) | $m$=1,113,978;<br>$h_{SNP}^2$=0.186(0.009)<br>intercept=1.070(0.025) |
| CD | No | $m$=1,148,525;<br>$h_{SNP}^2$=0.499(0.058)<br>intercept=1.023(0.011) | $m$=1,132,622;<br>$h_{SNP}^2$=0.501(0.057)<br>intercept=1.020(0.011) | $m$=1,115,890;<br>$h_{SNP}^2$=0.483 (0.058)<br>intercept=1.022 (0.011) |
| | Yes | $m$=1,147,392;<br>$h_{SNP}^2$=0.493(0.057)<br>intercept=1.022(0.011) | $m$=1,124,587;<br>$h_{SNP}^2$=0.503(0.056)<br>intercept=1.015(0.011) | $m$=1,112,341;<br>$h_{SNP}^2$=0.478 (0.057)<br>intercept=1.022 (0.011) |
| MDD | No | $m$=1,146,225;<br>$h_{SNP}^2$=0.059(0.003)<br>intercept=1.002(0.010) | $m$=1,130,372;<br>$h_{SNP}^2$=0.058(0.002)<br>intercept=0.998(0.010) | $m$=1,142,410;<br>$h_{SNP}^2$=0.056(0.002)<br>intercept=1.005(0.010) |
| | Yes | $m$=1,142,980;<br>$h_{SNP}^2$=0.059(0.002)<br>intercept=1.001(0.010) | $m$=1,120,468;<br>$h_{SNP}^2$=0.059(0.002)<br>intercept=0.993(0.010) | $m$=1,108,913;<br>$h_{SNP}^2$=0.056 (0.002)<br>intercept=1.003 (0.010) |
| BC | No | $m$=1,086,749;<br>$h_{SNP}^2$=0.129(0.012)<br>intercept=1.097(0.026) | $m$=1,106,705;<br>$h_{SNP}^2$=0.113(0.003)<br>intercept=0.932(0.016) | $m$=1,057,315;<br>$h_{SNP}^2$=0.1228 (0.012)<br>intercept=1.107 (0.030) |
| | Yes | $m$=1,085,928;<br>$h_{SNP}^2$=0.128(0.011)<br>intercept=1.091(0.023) | $m$=1,110,625;<br>$h_{SNP}^2$=0.113(0.003)<br>intercept=0.935(0.0163) | $m$=1,055,726;<br>$h_{SNP}^2$=0.123 (0.011)<br>intercept=1.086 (0.022) |
| OC | No | $m$=1,068,884;<br>$h_{SNP}^2$=0.050(0.010)<br>intercept=1.015(0.007) | $m$=1,054,645;<br>$h_{SNP}^2$=0.049(0.010)<br>intercept=1.016(0.007) | $m$=1,040,816;<br>$h_{SNP}^2$=0.047 (0.009)<br>intercept=1.018 (0.008) |
| | Yes | $m$=1,068,024;<br>$h_{SNP}^2$=0.050(0.010)<br>intercept=1.015(0.007) | $m$=1,052,337;<br>$h_{SNP}^2$=0.050(0.009)<br>intercept=1.014(0.007) | $m$=1,039,763;<br>$h_{SNP}^2$=0.047(0.009)<br>intercept=1.017(0.008) |

$m$: the number of variants; $h^2_{SNP}$: the SNP-based heritability estimate; intercept: LD score regression intercept. Values in parentheses are the stand errors of the corresponding estimates.

**Supplementary Table 16.** Computing resources used by DENTIST in the analyses of the UKB height data on chromosome 2.

| LD reference | Number of variants | Reference sample size | Memory usage | Number of CPUs | CPU time |
|---|---|---|---|---|---|
| **UKB.v3-20K** | 626K | 20,000 | <6G | 4 | 4h24min |
| **UKB.v3-20K** | 95K | 20,000 | <6G | 4 | 11min |
| **UKB.v3-8K** | 626K | 8,000 | <6G | 4 | 2h17min |
| **UKB.v3-8K** | 95K | 8,000 | <6G | 4 | 6min |

### Supplementary Notes

#### Supplementary Note 1. The exact and approximate models for z-statistic prediction

The method derived in the main text is under the null hypothesis of no association. Here we show a more general form of the derivation without this assumption. Following the definitions and notations in the main text, the exact form of the distribution of the z-statistic of a variant *i* in S1 conditioning on the variants in S2 is[1,2]

$$Z_i|\mathbf{Z}_t = \mathbf{z}_t \sim N(\mathbf{\Sigma}_{it}\mathbf{\Sigma}_{tt}^{-1}\mathbf{z}_t + (\mu_i - \mathbf{\Sigma}_{it}\mathbf{\Sigma}_{tt}^{-1}\boldsymbol{\mu}_t), \Sigma_{ii} - \mathbf{\Sigma}_{it}\mathbf{\Sigma}_{tt}^{-1}\mathbf{\Sigma}_{it}')$$

where $\mu_i = E(Z_i)$, $\boldsymbol{\mu}_t = \mathbf{Z}_t$ and all the other notations are the same as those in the main text. Compared to Equation 1 in the main text derived under the null model, this equation has an additional term $\mu_i - \mathbf{\Sigma}_{it}\mathbf{\Sigma}_{tt}^{-1}\boldsymbol{\mu}_t$, which is zero under the null hypothesis of no association. We demonstrate by numerical analysis that even under the alternative hypothesis, this term is small, and the z-statistic predicted based on the approximation method (derived under the null) is very close to that predicted based on the exact method (**Supplementary Figure 14**).

#### Supplementary Note 2. Acknowledgments

**UKB:** UK Biobank was established by the Wellcome Trust medical charity, Medical Research Council, Department of Health, Scottish Government and the Northwest Regional Development Agency. It has also had funding from the Welsh Assembly Government, British Heart Foundation and Diabetes UK.

**UK10K:** The UK10K project was funded by the Wellcome Trust award WT091310. Twins UK (TUK): TUK was funded by the Wellcome Trust and ENGAGE project grant agreement HEALTH-F4-2007–201413. The study also receives support from the Department of Health via the National Institute for Health Research (NIHR)-funded BioResource, Clinical Research Facility and Biomedical Research Centre based at Guy's and St. Thomas' NHS Foundation Trust in partnership with King's College London. Dr Spector is an NIHR senior Investigator and ERC Senior Researcher. Funding for the project was also provided by the British Heart Foundation grant PG/12/38/29615 (Dr Jamshidi). A full list of the investigators who contributed to the UK10K sequencing is available from http://www.UK10K.org.

**HRS:** HRS is supported by the National Institute on Aging (NIA U01AG009740). The genotyping was funded separately by the National Institute on Aging (RC2 AG036495, RC4 AG039029). Genotyping was conducted by the NIH Center for Inherited Disease Research (CIDR) at Johns Hopkins University. Genotyping quality control and final preparation of the data were performed by the Genetics Coordinating Center at the University of Washington.

**ARIC**: The Atherosclerosis Risk in Communities Study is carried out as a collaborative study supported by National Heart, Lung, and Blood Institute contracts (HHSN268201100005C, HHSN268201100006C, HHSN268201100007C, HHSN268201100008C, HHSN268201100009C, HHSN268201100010C, HHSN268201100011C, and HHSN268201100012C), R01HL087641, R01HL59367 and R01HL086694; National Human Genome Research Institute contract U01HG004402; and National Institutes of Health contract HHSN268200625226C. The authors thank the staff and participants of the ARIC study for their important contributions. Infrastructure was partly supported by Grant Number UL1RR025005, a component of the National Institutes of Health and NIH Roadmap for Medical Research.
